# Molecular basis of enhanced GLP-1 signaling mediated by GLP-1(9-36) in conjunction with LSN3318839

**DOI:** 10.1101/2024.07.21.604514

**Authors:** Jie Li, Guanyi Li, Yiting Mai, Xiao Liu, Dehua Yang, Qingtong Zhou, Ming-Wei Wang

## Abstract

Glucagon-like peptide-1 (GLP-1) receptor (GLP-1R) is essential for glucose metabolism and energy balance. It is primarily activated by the endogenous hormone GLP-1(7-36) and its weaker metabolite GLP-1(9-36). Although GLP-1(9-36) was previously considered biologically inactive, emerging evidence indicate that it exerts cardioprotective and insulinotropic actions. Here, we present cryo-electron microscopy structures showing that GLP-1(9-36) binds to an atypical site composed of the upper halves of transmembrane helices 1 (TM1) and 2 (TM2), extracellular loop 1, and the extracellular domain, distinct from the binding site of GLP-1(7-36). Meanwhile, LSN3318839, an orally effective positive allosteric modulator selective for GLP-1(9-36), anchors in the TM1-TM2 cleft and repositions GLP-1(9-36) to adopt a binding pose like GLP-1(7-36), thereby enhancing GLP-1R-mediated signal transduction. Mutagenesis and molecular dynamics simulations confirmed the crucial interactions and structural adjustments involved in this modulation. Our study uncovers a novel binding mode of GLP-1 metabolite and allosteric modulation that augments GLP-1R signaling, both are potentially insightful to unveil the physiological role of GLP-1(9-36) and design better small molecule therapeutics targeting GLP-1R.

## Introduction

Glucagon-like peptide 1 (GLP-1) receptor (GLP-1R) is a well-established drug target for type 2 diabetes and obesity. It is activated by the endogenous hormone GLP-1(7-36)NH_2_ [GLP-1(7-36) hereafter] to maintain glucose homeostasis^1^. GLP-1(7-36), produced through the proteolytic processing of proglucagon and released by L cells in the small intestine and colon, is rapidly degraded by dipeptidyl peptidase 4 with a half-life of approximately 2 minutes. The resulting metabolite, GLP-1(9-36)NH_2_ [GLP-1(9-36) hereafter], lacks the N-terminal dipeptide His7 and Ala8 (**Fig. S1A**)^2^ and constitutes approximately 60% of circulating GLP-1^3^. Compared to GLP-1(7-36), GLP-1(9-36) is a low potency and weak partial agonist of GLP-1R (**Fig. 1A**)^4^. It was initially believed to be biologically inactive. However, GLP-1(9-36) has been implicated in hepatic glucose production^5^, glycemic control^6^ and glucagonostatic actions^7,8^. Its cardiovascular^9,10^ and neuroprotective benefits^11,12^ were also reported.

**Figure 1.**
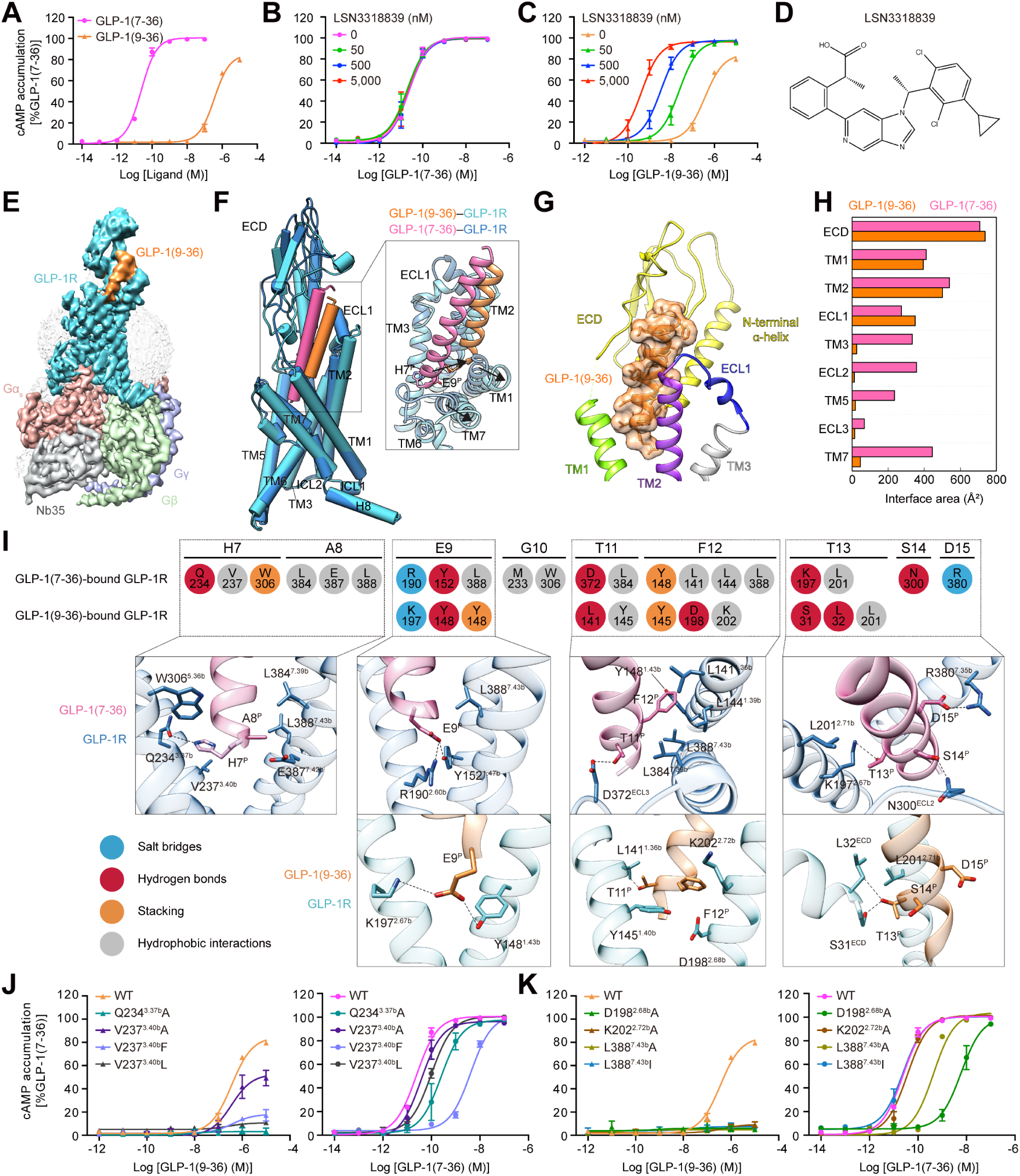
Molecular recognition of GLP-1(9-36) by GLP-1R. (A) Concentration–response curves of cAMP accumulation elicited by GLP-1(7-36) and GLP-1(9-36) at GLP-1R. (C-E) Effects of varying concentrations of LSN3318839 on GLP-1R-mediated cAMP accumulation induced by GLP-1(7-36) (B) and GLP-1(9-36) (C). Chemical structure of LSN3318839 (D). Data shown are means ± S.E.M. of at least three independent experiments performed in quadruplicate. (E) Cryo-EM density map of the GLP-1(9-36)–GLP-1R–G_s_ complex. (F) Superimposition of GLP-1(9-36)-bound and GLP-1(7-36)-bound GLP-1R structures. (G) Close-up view of the GLP-1(9-36)-binding pocket in GLP-1R. (H) Comparison of the interface area between individual segments (ECD, TMs and ECLs) of GLP-1R and GLP-1(9-36) or GLP-1(7-36). The interface areas were calculated using freeSASA. (I) Comparison of GLP-1R interactions with the N-terminal halves of GLP-1(9-36) and GLP-1(7-36), described by fingerprint strings encoding different interaction types of the surrounding residues. Close-up views of these interactions are shown with polar contacts depicted as black dashed lines. Receptor residues are labeled with class B1 GPCR numbering and colored by interaction type: sky blue (salt bridge), red (hydrogen bond), orange (stacking) and gray (hydrophobic interactions). (J, K) Effects of receptor mutations on GLP-1(9-36)- and GLP-1(7-36)-induced cAMP accumulation. Data shown are means ± S.E.M. of at least three independent experiments performed in quadruplicate.

LSN3318839 is an orally effective positive allosteric modulator selective for GLP-1(9-36). It transformed GLP-1(9-36) from a partial agonist (EC_50_ = 363 nM) to a full agonist (EC_50_ = 23 nM) at a concentration of 50 nM in cAMP accumulation assays (**Fig. 1B-D**)^13^. Here, we report the cryo-electron microscopy (cryo-EM) structures of GLP-1(9-36)-bound GLP-1R in complex with G_s_ alone and in the presence of LSN3318839, with global resolutions of 3.0 Å and 3.2 Å, respectively (**Figs. 1, 2, S1-S3 and Table S1**). Together with mutagenesis and molecular dynamics (MD) simulations, we propose a peptide-binding mechanism previously unseen in GLP-1R and provide a structural basis of enhanced GLP-1R signaling allosterically modulated by LSN3318839.

**Figure 2.**
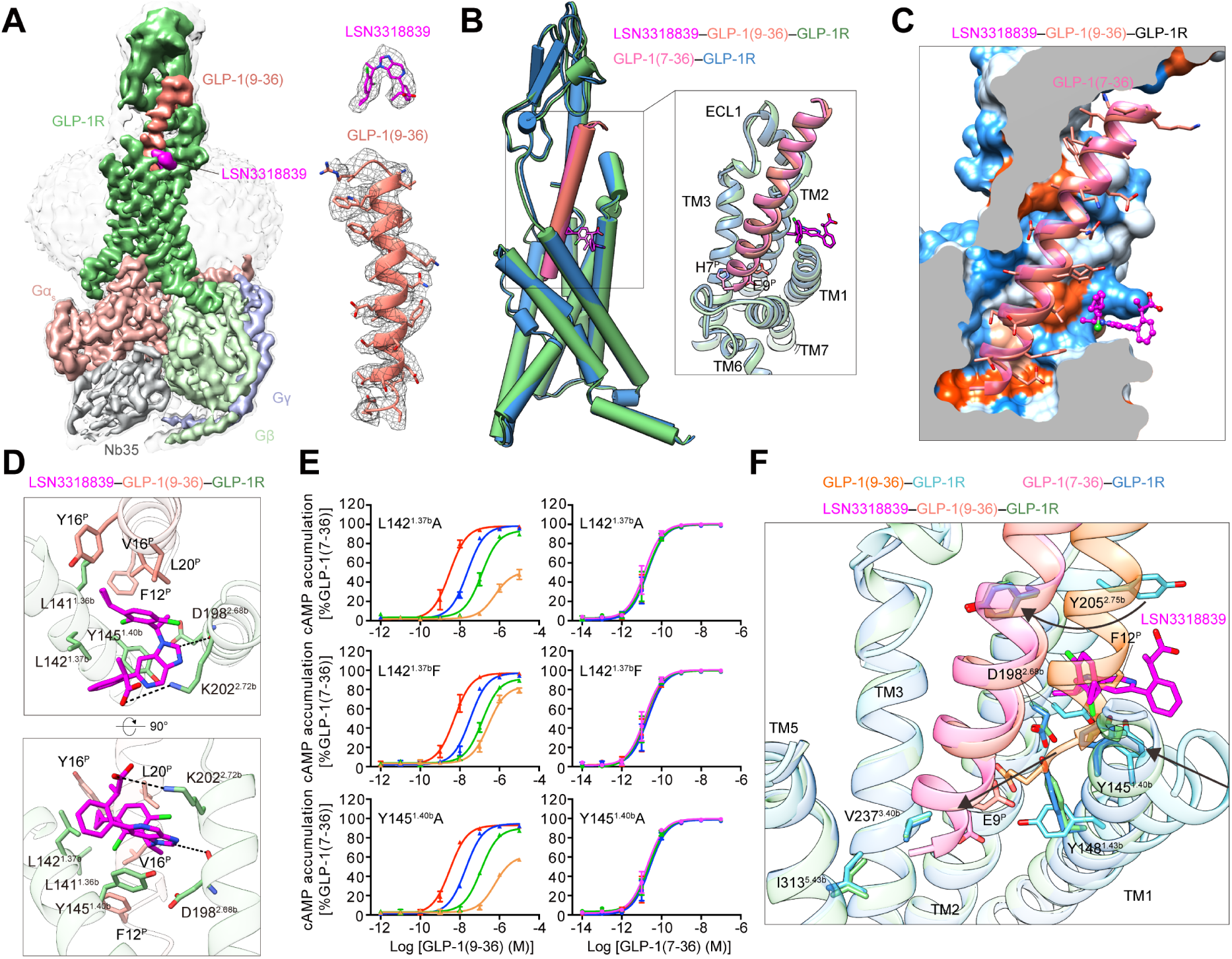
Allosteric modulation of LSN3318839 at GLP-1(9-36)–GLP-1R. (A) Cryo-EM density maps of the LSN3318839–GLP-1(9-36)–GLP-1R–G_s_ complexes. Atomic models and EM densities of the bound ligands are shown as sticks and surfaces, respectively. (B) Superimposition of LSN3318839–GLP-1(9-36)–GLP-1R and GLP-1(7-36)–GLP-1R structures. A close-up view highlights that LSN3318839 induces GLP-1(9-36) to adopt a conformation similar to GLP-1(7-36). (C) Surface representation of the peptide-binding pocket in GLP-1R bound by LSN3318839 and GLP-1(9-36). GLP-1(9-36) is shown as ribbon and stick. Receptor surfaces are colored from dodger blue for the most hydrophilic regions, to white, to orange red for the most hydrophobic regions. (D) Interactions between LSN3318839 and GLP-1(9-36)–GLP-1R. Residues involved in LSN3318839 recognition are shown as sticks. Polar contacts are shown as black dashed lines. (E) Effects of receptor mutations on cAMP accumulation elicited by GLP-1(9-36) (left) and GLP-1(7-36) (right) in the presence of LSN3318839. Data shown are means ± S.E.M. of at least three independent experiments performed in quadruplicate. (F) A close-up view of interactions between peptide and TMD pocket for superimposed GLP-1(9-36)–GLP-1R, LSN3318839–GLP-1(9-36)–GLP-1R and GLP-1(7-36)–GLP-1R structures. Arrows indicate conformational transition of GLP-1(9-36)– GLP-1R upon LSN3318839 binding.

## Results

### GLP-1(9-36) binding

Superimposing the structures of GLP-1(9-36)–GLP-1R and GLP-1(7-36)–GLP-1R^14^ reveals that GLP-1(9-36) and GLP-1(7-36) interacts with two distinct sites on GLP-1R, despite both adopting a single continuous helix (**Fig. 1F**). The binding site for GLP-1(9-36) is formed by the upper halves of transmembrane helices 1 (TM1) and 2 (TM2), extracellular loop 1 (ECL1) and the extracellular domain (ECD), with negligible involvement of the residues in TM3, ECL2, TM5, ECL3 and TM7 commonly participate in the hormone recognition of class B1 G protein-coupled receptors (**Figs. 1G-I and S4**). Compared to GLP-1(7-36), the N-terminal, middle and C-terminal segments of GLP-1(9-36) shift outward by 7.3 Å, 12.2 Å and 10.3 Å, measured at the Cα atoms of E9^P^ (P indicates that the residue belongs to the peptide), S18^P^ and E27^P^, respectively. This movement renders the upper halves of TM1, TM2 and TM7 outward by 6.0 Å, 4.9 Å and 6.6 Å, measured at the Cα atoms of E139^1.34b^, K202^2.72b^ and T378^7.33b^ (class B1 GPCR numbering in superscript), respectively (**Fig. 1F**). Upon GLP-1(9-36) binding, the most significant conformational changes occur in ECL1 and ECD, compared to the GLP-1(7-36)-bound GLP-1R (**Fig. 1F**). ECL1, which forms a robust α-helix adjacent to TM2 in most peptide-bound GLP-1R structures, makes a short α-helix closer to TM3. Meanwhile, the ECD rotates downward to fill the transmembrane domain (TMD) and accommodate GLP-1(9-36), evidenced by a 10.5 Å downward movement of the N-terminal α-helix (measured at the Cα atom of S31). These conformational features created a well-defined GLP-1R pocket for GLP-1(9-36) with a total interface area of 1,891 Å^2^, significantly lower than that of GLP-1(7-36) (3,316 Å^2^), consistent with its weak potency (**Fig. 1A**). TMs 3-6 and the intracellular halves of TM1, TM2 and TM7 of the GLP-1(9-36)-bound GLP-1R align well with that bound by GLP-1(7-36), indicating a conserved TMD architecture (**Fig. 1F**).

### Receptor interaction

Interaction of GLP-1(9-36) with GLP-1R involves a complex network of polar and nonpolar nature, utilizing different residues compared to GLP-1(7-36) (**Figs. 1I, S4D and Table S2**). E9^P^, the first N-terminal residue of GLP-1(9-36), forms a salt bridge with K197^2.67b^ and a hydrogen bond with Y148^1.43b^. In contrast, these two interactions are with R190^2.60b^ and Y152^1.47b^, respectively, in the case of GLP-1(7-36) (**Fig. 1I**). The first two N-terminal residues of GLP-1(7-36), which GLP-1(9-36) lacks, H7^P^ and A8^P^, are located above the central polar network and exhibit extensive interactions with TMs 3, 5 and 7, including Q234^3.37b^ and V237^3.40b^. Such interactions are absent in the case of GLP-1(9-36). Mutations Q234^3.37b^A, V237^3.40b^L and V237^3.40b^F decreased the potency (EC_50_) of GLP-1(7-36) by 15-fold, 6-fold and 257-fold, respectively (**Fig. 1J, K and Tables S3 and S4**). However, their impact on cAMP signaling induced by 10 μM GLP-1(9-36) was different: from 85% in wildtype (WT) GLP-1R to 3% (Q234^3.37b^A), 10% (V237^3.40b^L) and 18% (V237^3.40b^F), relative to the maximum response (100%) induced by GLP-1(7-36) (**Fig. 1J**), indicating that these three residues are essential for GLP-1(7-36).

T11^P^ in GLP-1(9-36) primarily interacts with TM1, forming a hydrogen bond with L141^1.36b^ and hydrophobic contacts with Y145^1.40b^, while T13^P^ interacts with the ECD by making two hydrogen bonds with S31^ECD^ and L32^ECD^. In GLP-1(7-36), T11^P^ and T13^P^ mainly interact with ECL3/TM7 and TM2, respectively (**Fig. 1I**). F12^P^ in GLP-1(9-36) is positioned in the TM1-TM2 cleft, stacking with Y145^1.40b^, forming an aromatic hydrogen bond with D198^2.68b^, and making hydrophobic contacts with K202^2.72b^. For GLP-1(7-36), F12^P^ stacks with Y148^1.43b^ and has hydrophobic contacts with L141^1.36b^, L144^1.39b^ and L388^7.43b^ (**Fig. 1I**). The critical roles of these residues are highlighted by mutagenesis studies, where D198^2.68b^A, K202^2.72b^A, L388^7.43b^A and L388^7.43b^I abolished GLP-1(9-36)-induced cAMP accumulation, but still allowed GLP-1(7-36) signaling (**Fig. 1K**).

The C-terminal half of GLP-1(9-36) is recognized by the ECD, ECL1 and TM2 (**Fig. S4**). Three residues (V16^P^, Y19^P^ and L20^P^) engage in stacking and hydrophobic contacts with the sidechains of TM2 and ECL1 residues including L201^2.71b^, Y205^2.75b^ and A208^ECL1^. ECL1 forms two hydrogen bonds with Q23^P^ (via W214^ECL1^) and E27^P^ (via Q211^ECL1^), as well as stacking interaction with F28^P^ (via W214^ECL1^). The ECD recognizes GLP-1(9-36) by forming multiple hydrogen bonds (via S17^P^, A24^P^ and I29^P^) and hydrophobic contacts (via V16^P^, L20^P^, A25^P^ and I29^P^). These residues have similar interactions between the two peptides (**Table S2**).

### LSN3318839 binding

As shown in **Fig. 2B**, the LSN3318839–GLP-1(9-36)–GLP-1R complex closely resembles that of GLP-1(7-36)–GLP-1R, with a Cα root mean square deviation (RMSD) value of 0.64 Å, significantly different from the GLP-1(9-36)–GLP-1R structure (Cα RMSD of 1.65 Å). The bound peptide and surrounding TMs and ECLs of GLP-1R in the presence of both GLP-1(9-36) and LSN3318839 are well-aligned with those in the GLP-1(7-36)–GLP-1R complex (**Fig. 2A and 2B**). The interface area of GLP-1(9-36) with GLP-1R increases from 1,891 Å² [GLP-1(9-36) alone] to 3,010 Å² in the presence of LSN3318839, nearly matching that of GLP-1(7-36) (3,316 Å²). It appears that LSN3318839 induces GLP-1(9-36) to adopt a binding pose identical to that of GLP-1(7-36), supported by an enhanced cAMP signaling when LSN3318839 is present (**Fig. 2C**).

LSN3318839 anchors in the TM1-TM2 cleft by extensively interacting with residues in TM1, TM2 and GLP-1(9-36) (**Fig. 2B and 2D**). The central 5-azabenzimidazole group of LSN3318839 is lifted by the base of the binding pocket (Y145^1.40b^ and D198^2.68b^) through π–π stacking with Y145^1.40b^, cation–π stacking with K202^2.72b^, and an aromatic hydrogen bond with D198^2.68b^. The dichlorophenyl ring and cyclopropane, which closely pack with GLP-1(9-36), occupy a hydrophobic pocket and receive extensive hydrophobic interactions with L141^1.36b^, L142^1.37b^, L201^2.71b^, F12^P^, V16^P^, Y19^P^ and L20^P^. The external 2-phenylpropionate moiety is stabilized by stacking with Y145^1.40b^ and hydrophobic interactions with L142^1.37b^, while the carboxylic acid forms a salt bridge with K202^2.72b^. Mutagenesis studies show that 5 μM LSN3318839 increased the potency of GLP-1(9-36) by 891-fold (EC_50_ shifted from 0.36 μM to 0.41 nM) for WT GLP-1R. However, this enhancement was significantly impaired by mutations such as L142^1.37b^A, Y145^1.40b^A and L142^1.37b^F (decreasing to 200-fold, 186-fold and 52-fold, respectively) (**Fig. 2E**). Importantly, these mutations did not affect the potency of GLP-1(7-36), verifying the selectivity of LSN3318839 for GLP-1(9-36) (**Fig. 2E**). In the MD simulations, LSN3318839 steadily locked into the TM1-TM2 cleft, exhibiting binding poses similar to that observed in the cryo-EM structure (**Fig. S5**).

### Allosteric modulation

Comparing the structures of GLP-1(9-36)–GLP-1R with and without LSN3318839 reveals significant spatial overlaps between the first N-terminal helical turn of GLP-1(9-36) (residues T11^P^–S14^P^) and LSN3318839, with F12^P^ being the most prominent overlay (**Fig. 2F**). This suggests that LSN3318839 binding likely pushes the GLP-1(9-36) N terminus back toward the TMD core, where GLP-1(7-36) locates. Such repositioning is further promoted by three microswitches: Y148^1.43b^, D198^2.68b^ and Y205^2.75b^ (**Fig. 2F**). In the absence of LSN3318839, the N-terminal tip of GLP-1(9-36) is stabilized by the downward orientation of Y148^1.43b^ as well as the Y145^1.40b–^D198^2.68b^ pair which tightens the TM1-TM2 cleft. The C-terminal half is stabilized by interactions with the ECD N terminus α-helix, TM2 and ECL1, with Y205^2.75b^ pointing outward to the TM1-TM2 cleft allowing the entrance of ECD. Upon LSN3318839 binding, Y148^1.43b^ is released to rotate upward forming a hydrogen bond with the inward-rotated D198^2.68b^, stably holding the peptide N terminus. Concurrently, Y205^2.75b^ shifts inward to approach ECL2, making a hydrogen bond with R299^ECL2^ and establishing a polar network involving Q221^ECL1^ and E21^P^ to stabilize binding. These coordinated movements triggered by LSN3318839 ensure that GLP-1(9-36) adopts a binding conformation similar to GLP-1(7-36). Consistently, in the MD simulation, the N terminus of GLP-1(9-36) exhibits stable interactions within the TMD, closely resembling those observed in the cryo-EM structure. However, removing LSN3318839 from the simulation system significantly disrupted the binding of the GLP-1(9-36) N terminus (**Fig. S6**).

## Discussion

Present study demonstrates that GLP-1(9-36) adopts a binding conformation in GLP-1R distinct from previously reported liganded GLP-1R structures. It engages GLP-1R through interactions with TM1, TM2, ECL1 and ECD, rather than TM3, ECL2, TM5, ECL3 and TM7 with GLP-1(7-36). This unique binding mode explains why the metabolite is almost inactive at GLP-1R. Nonetheless, this structural feature could be transformed to that of GLP-1(7-36) by a selective allosteric modulator LSN3318839, thereby enhancing GLP-1R-mediated signal transduction. Mutagenesis and MD simulations confirmed the crucial interactions and structural adjustments involved in this modulation. Our study uncovers a novel binding mode of GLP-1 metabolite and allosteric modulation that augments GLP-1R signaling, both are potentially insightful to unveil the physiological role of GLP-1(9-36) and design better small molecule therapeutics targeting GLP-1R.

## Materials and methods

### Cell culture

*Spodoptera frugiperda* (*Sf*9) insect cells (Expression Systems) were cultured in ESF 921 insect cell culture medium (Expression Systems) at 27°C and 120 rpm. HEK-293T cells were grown in Dulbecco’s modified Eagle’s medium (DMEM, Life Technologies) supplemented with 10% fetal bovine serum (FBS, Gibco) and maintained in a humidified chamber supplemented with 5% CO_2_ at 37°C.

### Constructs

The native signal sequence (M1-P23) of human GLP-1R was replaced by the haemagglutinin (HA) signal peptide to facilitate receptor expression. To obtain the GLP-1R–G_s_ complex with good stability and homogeneity, the NanoBiT tethering strategy was used^15^, in which the C terminus of GLP-1R was directly attached to the LgBiT subunit^16^ (Promega), followed by a tobacco etch virus (TEV) protease cleavage site and a double maltose binding protein (MBP) tag (Fig. S1A). The C terminus of rat Gβ1 was linked to peptide 86 subunit^16,17^ (also named as HiBiT, Promega) with a 15-amino acid polypeptide (GSSGGGGSGGGGSSG) linker. An engineered G_s_ construct (G112)^18^ was used for expression and purification of the complexes. The constructs were cloned into both pcDNA3.1 and pFastBac vectors for functional assays in mammalian cells and protein expression in insect cells, respectively. These modifications of the receptor had no effect on ligand binding and receptor activation^15,19^.

### Expression and purification

The Bac-to-Bac Baculovirus Expression System (Invitrogen) was used to generate high titer recombinant baculovirus for GLP-1R-LgBiT-2MBP, Gα_s_, Gβ1-HiBiT and Gγ2. P0 viral stock was produced by transfecting 5 μg recombinant bacmids into *Sf9* cells (2.5 mL, density of 1 million cells per mL) for 96 h incubation and then used to produce P1 and P2 baculovirus. The GLP-1R-LgBiT-2MBP, G112, Gβ1-HiBiT and Gγ2 were co-expressed at multiplicity of infection (MOI) ratio of 1:1:1:1 by infecting *Sf9* cells at a density of 3 million cells per mL with P2 baculovirus (viral titers > 90%). The culture was harvested by centrifugation after 48 h post-infection, and cell pellets were stored at −80°C until use.

The cell pellets were thawed and lysed in a buffer containing 20 mM HEPES, pH 7.5, 100 mM NaCl, 10% (v/v) glycerol, 10 mM MgCl_2_, 1 mM MnCl_2_ and 100 μM TCEP (Sigma-Aldrich) supplemented with EDTA-free protease inhibitor cocktail (TragetMol) by dounce homogenization. The GLP-1(9-36)–GLP-1R–G_s_ complex formation was initiated by the addition of 100 μM GLP-1(9-36), 10 μg/mL nanobody 35 (Nb35) and 25 mU/mL apyrase (New England Bio-Labs). As for the LSN3318839–GLP-1(9-36)–GLP-1R–G_s_ complex formation, 10 μM GLP-1(9-36), 50 μM LSN3318839, 10 μg/mL Nb35 and 25 mU/mL apyrase (New England Bio-Labs) were added.

After 1.5 h incubation at room temperature (RT), the membrane was solubilized in the buffer above supplemented with 0.5% (w/v) lauryl maltose neopentyl glycol (LMNG, Anatrace) and 0.1% (w/v) cholesterol hemisuccinate (CHS, Anatrace) for 2 h at 4°C. The supernatant was isolated by centrifugation at 65,000× *g* for 30 min and incubated with amylose resin (New England Bio-Labs) for 2 h at 4°C. The resin was then collected by centrifugation at 600× *g* for 10 min and washed in gravity flow column (Sangon Biotech) with five column volumes of buffer (pH 7.5) containing 20 mM HEPES, 100 mM NaCl, 10% (v/v) glycerol, 5 mM MgCl_2_, 1 mM MnCl_2_, 25 μM TCEP, 0.1% (w/v) LMNG, 0.02% (w/v) CHS and 10 μM ligand, followed by washing with fifteen column volumes of buffer (pH 7.5) containing 20 mM HEPES, 100 mM NaCl, 10% (v/v) glycerol, 5 mM MgCl_2_, 1 mM MnCl_2_, 25 μM TCEP, 0.03% (w/v) LMNG, 0.01% (w/v) glyco-diosgenin (GDN, Anatrace), 0.008% (w/v) CHS and 10 μM ligand. The protein was then incubated overnight with TEV protease on the column to remove the C-terminal 2MBP-tag in the buffer above at 4°C. The flow-through was collected the next day and concentrated with a Amicon Ultra centrifugal filter (molecular weight cut-off of 100 kDa, Millipore) and then subjected to a Superdex 200 Increase 10/300GL column (Cytiva) with running buffer (pH 7.5) containing 20 mM HEPES, 100 mM NaCl, 10 mM MgCl_2_, 100 μM TCEP, 10 μM ligand, 0.00075% (w/v) LMNG, 0.00025% (w/v) GDN and 0.0002% (w/v) CHS. The fractions of the monomeric complex were collected and concentrated to 11 mg/mL for electron microscopy examination.

### Expression and purification of Nb35

Nb35 with a C-terminal 6× His-tag was expressed in the periplasm of E. *coli* BL21 (DE3), extracted, and purified by nickel affinity chromatography as previously described^20^. Briefly, the Nb35 target gene was transformed in BL21 (DE3) and grown in a TB culture medium with 100 μg/mL ampicillin, 2 mM MgCl_2_ and 0.1% (w/v) glucose at 37°C, 180 rpm. Then the expression was induced by adding 1 mM IPTG when OD_600_ reached 0.7–1.2. The cell pellet was collected by centrifugation after overnight incubation at 28°C, 180 rpm and stored at −80°C until use. The HiLoad 16/600 Superdex 75 column (GE Healthcare) was used to separate the monomeric fractions of Nb35 with running buffer (pH 7.5) containing 20 mM HEPES and 100 mM NaCl. The purified Nb35 containing 30% (v/v) glycerol was flash-frozen by liquid nitrogen and stored at −80°C until use.

### Structure determination

To prepare a high-quality GLP-1(9-36)–GLP-1R–G_s_ complex, the NanoBiT tethering strategy was employed (**Fig. S1B**)^16,21^. These protein complexes were subsequently purified as monodispersed peaks on size exclusion chromatography (SEC) demonstrating a good homogeneity and verified by SDS gel to confirm all the expected components (**Fig. S1C and S1D**). After sample preparation, cryo-EM data collection, single particle analysis and three-dimensional (3D) classification were performed that yielded consensus density maps with overall resolutions of 3.0 Å and 3.2 Å for the GLP-1(9-36)–GLP-1R–G_s_ and LSN3318839– GLP-1(9-36)–GLP-1R–G_s_ complexes, respectively (**Figs. 1E, 2A, S2, S3 and Table S1**). The cryo-EM maps enabled near-atomic modeling for most regions of the complexes, except the stalk between transmembrane helix 1 (TM1) and the extracellular domain (ECD) of GLP-1R (**Fig. S3**). The presence of GLP-1(9-36), especially with LSN3318839, was clear enough in the two cryo-EM maps that enabled structural modeling of the backbones and sidechains of most residues (**Figs. 1A and S3**). Additionally, residues L339 to L343 in the intracellular loop 3 (ICL3) of the LSN3318839–GLP-1(9-36)–GLP-1R–G_s_ complex, residues F369 to E373 in the extracellular loop 3 (ECL3) of the GLP-1(9-36)–GLP-1R–G_s_ complex, and residues K130 to S135 in the GLP-1R ECD in these two complexes were poorly resolved and not modeled in the structures.

#### Cryo-EM data acquisition

The purified sample (3 μL) was applied to glow-discharged holey carbon grids (Quantifoil R1.2/1.3, 300 mesh), and subsequently vitrified using a Vitrobot Mark IV (ThermoFisher Scientific) set at 100% humidity and 4°C. Cryo-EM images were collected on a Titan Krios microscope (FEI) equipped with a Gatan energy filter and K3 direct electron detector and performed using serialEM. The microscope was operated at 300 kV accelerating voltage and a calibrated magnification of ×81,000 corresponding to a pixel size of 1.071 Å. An accumulated dose of 80 electrons per Å^2^ was fractionated into a movie stack of 36 frames, with a defocus range of −1.2 to −2.2 μm. Totally, 11,396 images for the GLP-1(9-36)–GLP-1R–G_s_ complex and 6,175 images for the LSN3318839–GLP-1(9-36)–GLP-1R–G_s_ complex were collected.

#### Cryo-EM data processing

For the LSN3318839–GLP-1(9-36)–GLP-1R–G_s_ complex, the micrographs were processed using cryoSPARC v.4.4.121 with patch motion correction to restore the correct information of the original cryo-EM images. The contrast transfer function (CTF) parameters were estimated using patch CTF estimation. Template picking referenced from the cryo-EM map of the compound 2−GLP-1R−G_s_ complex (EMDB code: EMD-30867) yielded 6,146,299 particle projections, which were subjected to two rounds of 2D classification to discard false positive particles or particles categorized in poorly defined classes, producing 1,497,042 particle projections. The *ab-initio* reconstruction and hetero refinement were applied to divide particles into five subsets, a selected subset of 662,976 particle projections was subjected to further processing. To improve the density quality of specific regions, this subset of particle projections was subjected to 3D auto-refinement with a mask on the complex, which was subsequently subjected to another round of 3D classifications with a mask on the ECD. After the last round of NU refinement, the final map of the LSN3318839–GLP-1(9-36)–GLP-1R–G_s_ complex has an indicated global resolution of 3.22 Å by the 0.143 criteria of the gold-standard Fourier shell correlation (FSC), based on a dataset of 238,182 particles.

For the GLP-1(9-36)–GLP-1R–G_s_ complex, 11,396 movies were motion corrected and CTF estimated, and 12,688,111 particles were auto-picked by template picker referenced to the LSN3318839–GLP-1(9-36)–GLP-1R–G_s_ map we obtained. After two rounds of 2D classification, four rounds of 3D classification and hetero refinement, and one round of non-uniform refinement and local refinement, 99,120 particles were used to generate a map with an indicated global resolution of 3.00 Å at an FSC of 0.143.

#### Model building and refinement

The cryo-EM structures of the and GLP-1(7-36)–GLP-1R–G_s_ (PDB code: 6X18)^14^ and LSN3318839–GLP-1(7-36)–GLP-1R–G_s_ (PDB code: 6VCB)^22^ were employed as the initial templates for modeling the GLP-1(9-36)–GLP-1R–G_s_ and LSN3318839–GLP-1(9-36)–GLP-1R–G_s_ complexes, respectively. The initial models were fitted into the EM density map using the UCSF Chimera 1.17.3^23,24^, followed by interactive rounds of manual adjustment and automated refinement in COOT 0.9.8.92^25^ and Phenix 1.21^26^, respectively. The final refinement statistics were validated using the module comprehensive validation (cryo-EM) in Phenix 1.21. Structural figures were prepared in UCSF Chimera 1.17.3, UCSF ChimeraX v1.0 and PyMOL 2.5.8 (https://pymol.org/2/). The final refinement statistics are provided in Table S1.

### cAMP accumulation assay

Small molecule or peptide stimulated cAMP accumulation was measured by a LANCE Ultra cAMP kit (PerkinElmer). Briefly, following 24 h transfection with various constructs, HEK-293T cells were washed once with Dulbecco’s phosphate-buffered saline (DPBS) and digested by 0.2% (w/v) EDTA in DPBS. After centrifugation, cell pellets were then collected and resuspended with stimulation buffer [HBSS supplemented with 5 mM HEPES, 0.5 mM IBMX and 0.1% (w/v) BSA, pH 7.4] to a density of 0.6 million cells per mL and added to 384-well culture plates (3,000 cells per well). Cells were stimulated by serial concentrations of ligand for 40 min at RT. The reaction was terminated by adding 5 μL Eu-cAMP tracer and 5 μL ULight-anti-cAMP. After 1 h incubation at RT, cAMP signals were detected by measuring the fluorescence intensity at 620 nm and 665 nm via an EnVision multilabel plate reader (PerkinElmer).

### Molecular dynamics simulation

Molecular dynamic (MD) simulations were performed by Gromacs 2021.4^27^. The MD simulation of the LSN3318839–GLP-1(9-36)–GLP-1R complex was built based on the cryo-EM structure of the LSN3318839–GLP-1(9-36)–GLP-1R–G_s_ complex and prepared by the Protein Preparation Wizard (Schrödinger 2017-4) with the G protein and Nb35 removed. The receptor chain termini were capped with acetyl and methylamide. All titratable residues were left in their dominant state at pH 7.0. To build MD simulation systems, the complexes were embedded in a bilayer composed of 257 POPC lipids and solvated with 0.15 M NaCl in explicit TIP3P waters using CHARMM-GUI Membrane Builder v3.5^28^. The CHARMM36m force field^29^ was adopted for protein, peptides, lipids and salt ions. The parameter of LSN3318839 was generated using the CHARMM General Force Field (CGenFF)^30^. The Particle Mesh Ewald (PME) method was used to treat all electrostatic interactions beyond a cut-off of 12 Å and the bonds involving hydrogen atoms were constrained using LINCS algorithm^31^. The complex system was first relaxed using the steepest descent energy minimization, followed by slow heating of the system to 310 K with restraints. The restraints were reduced gradually over 10 ns. Finally, restrain-free production run was carried out for each simulation, with a time step of 2 fs in the NPT ensemble at 310 K and 1 bar using the Nose-Hoover thermostat and the semi-isotropic Parrinello-Rahman barostat^32^, respectively. The interface area was calculated by the program FreeSASA 2.0, using the Sharke-Rupley algorithm with a probe radius of 1.2 Å^33^. Similar simulation procedure and analysis were adopted for the MD simulations of GLP-1(9-36)–GLP-1R complex by removing LSN3318839.

### Statistical analysis

All functional study data were analyzed using Prism 10 (GraphPad) and presented as means ± S.E.M. from at least three independent experiments. Concentration-response curves were evaluated with a three-parameter logistic equation. The significance was determined with either two-tailed Student’s t-test or one-way ANOVA, and *P* < 0.05 was considered statistically significant.

### Data availability

All relevant data are available from the authors and/or included in the manuscript or Supplementary Information. The atomic coordinates have been deposited in the Protein Data Bank (PDB) under accession codes: XXXX [GLP-1(9-36)–GLP-1R–G_s_] and XXXX [LSN3318839–GLP-1(9-36)–GLP-1R–G_s_], and the electron microscopy maps have been deposited in the Electron Microscopy Data Bank (EMDB) under accession codes: EMD-XXXXX [GLP-1(9-36)–GLP-1R–G_s_] and EMD-XXXXX [LSN3318839–GLP-1(9-36)–GLP-1R–G_s_].

## Acknowledgements

We are grateful to Wenxin Li, Wenbo Feng, Chenyu Ye, Yingna Xu, Yan Chen, Xianyue Chen, Jiahui Yan and Qiuying Wang for technical assistance. The cryo-EM data were collected at the Cryo-Electron Microscopy Research Center, Shanghai Institute of Materia Medica, Chinese Academy of Sciences. This work was partially supported by the National Natural Science Foundation of China 82273961 (M.-W.W.), 82073904 (M.-W.W.) and 81872915 (M.-W.W.); STI2030-Major Project 2021ZD0203400 (Q.T.Z.); and Hainan Provincial Major Science and Technology Project ZDKJ2021028 (D.H.Y. and Q.T.Z.).

## Author contributions

J.L., G.Y.L. and Q.T.Z. performed research; Y.T.M. and X.L. assisted in protein purification and data collection; J.L., Q.T.Z., D.H.Y. and M.-W.W. analyzed the data and wrote the manuscript with inputs from all co-authors; M.-W.W. initiated the project and supervised the studies.

---

**Figure S1.**
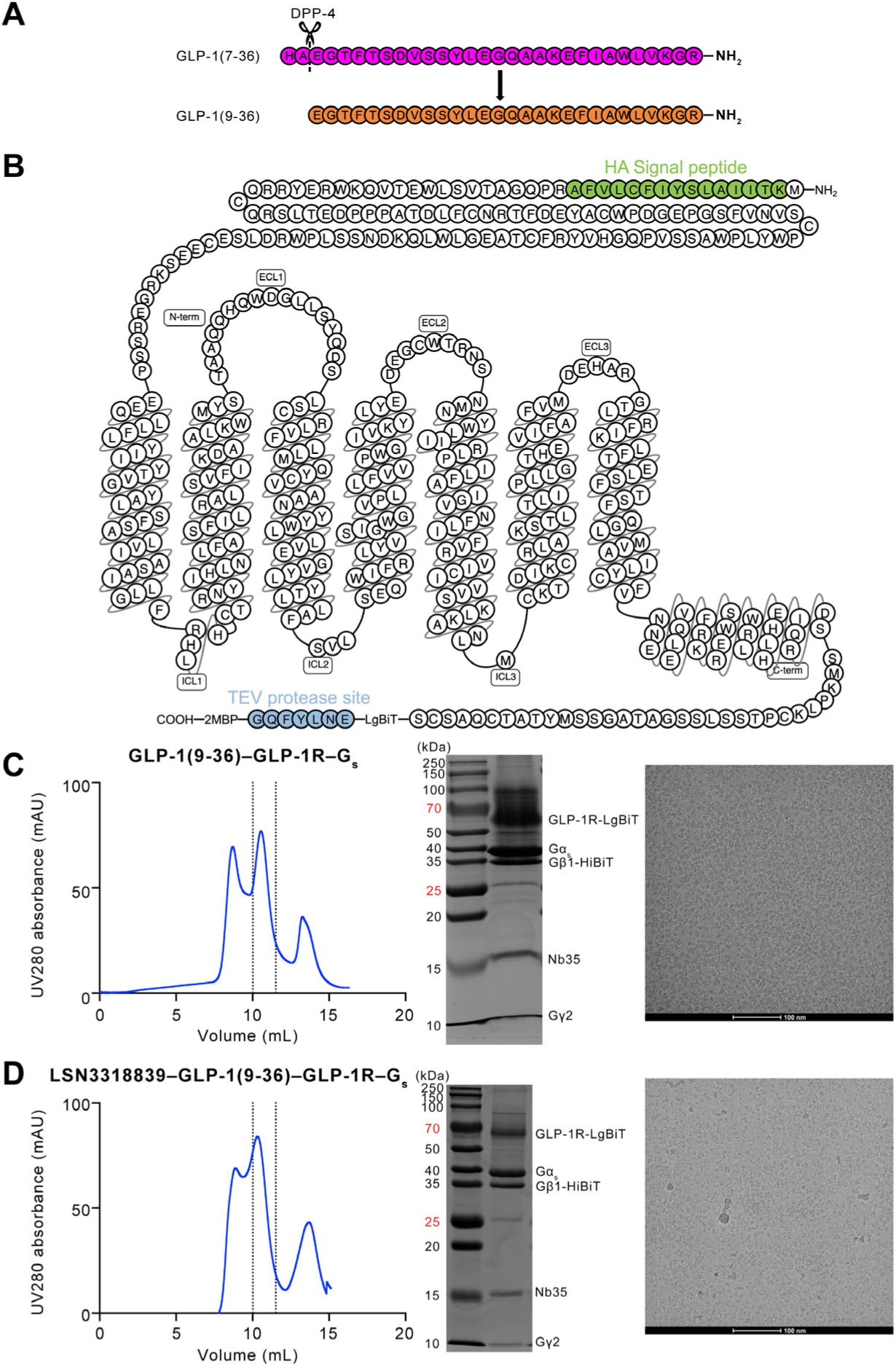
Purification and cryo-EM image of the GLP-1(9-36)–GLP-1R–G_s_ and LSN3318839–GLP-1(9-36)–GLP-1R–G_s_ complexes. (A) GLP-1(9-36) is a major metabolite of GLP-1(7-36) formed by dipeptidyl peptidase 4 (DPP-4) cleavage, truncating the N-terminal dipeptide His7-Ala8 from GLP-1(7-36). (B) Schematic diagram of the GLP-1R construct used for the structure determination. (C, D) Analytical size-exclusion chromatography (left), SDS-PAGE/Coomassie blue stain (middle) and representative cryo-EM micrograph (right, scale bar: 100 nm) of the purified GLP-1(9-36)–GLP-1R–G_s_ (C) and LSN3318839-GLP-1(9-36)–GLP-1R–G_s_ (D) complexes.

**Figure S2.**
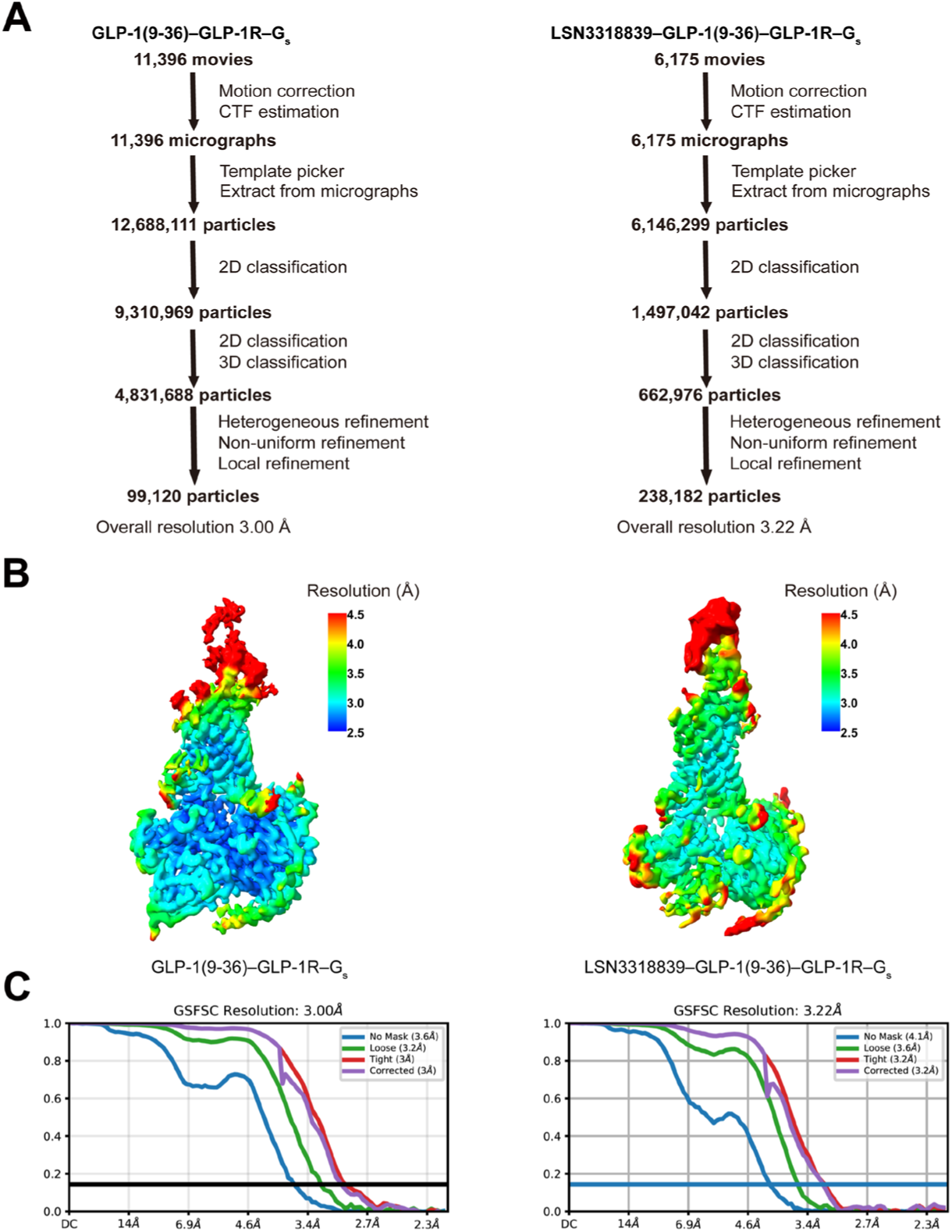
Cryo-EM data processing and validation of the GLP-1(9-36)–GLP-1R–G_s_ and LSN3318839–GLP-1(9-36)–GLP-1R–G_s_ complexes. (A) Cryo-EM data processing flow charts for the GLP-1(9-36)–GLP-1R–G_s_ (left) and LSN3318839–GLP-1(9-36)–GLP-1R–G_s_ (right) complexes. (B) Density maps colored by local resolution for the GLP-1(9-36)–GLP-1R–G_s_ (left) and LSN3318839–GLP-1(9-36)–GLP-1R–G_s_ (right) complexes. (C) Gold standard Fourier shell correlation (FSC) curves of overall refined structures, indicating the global resolution at 0.143 FSC threshold.

**Figure S3.**
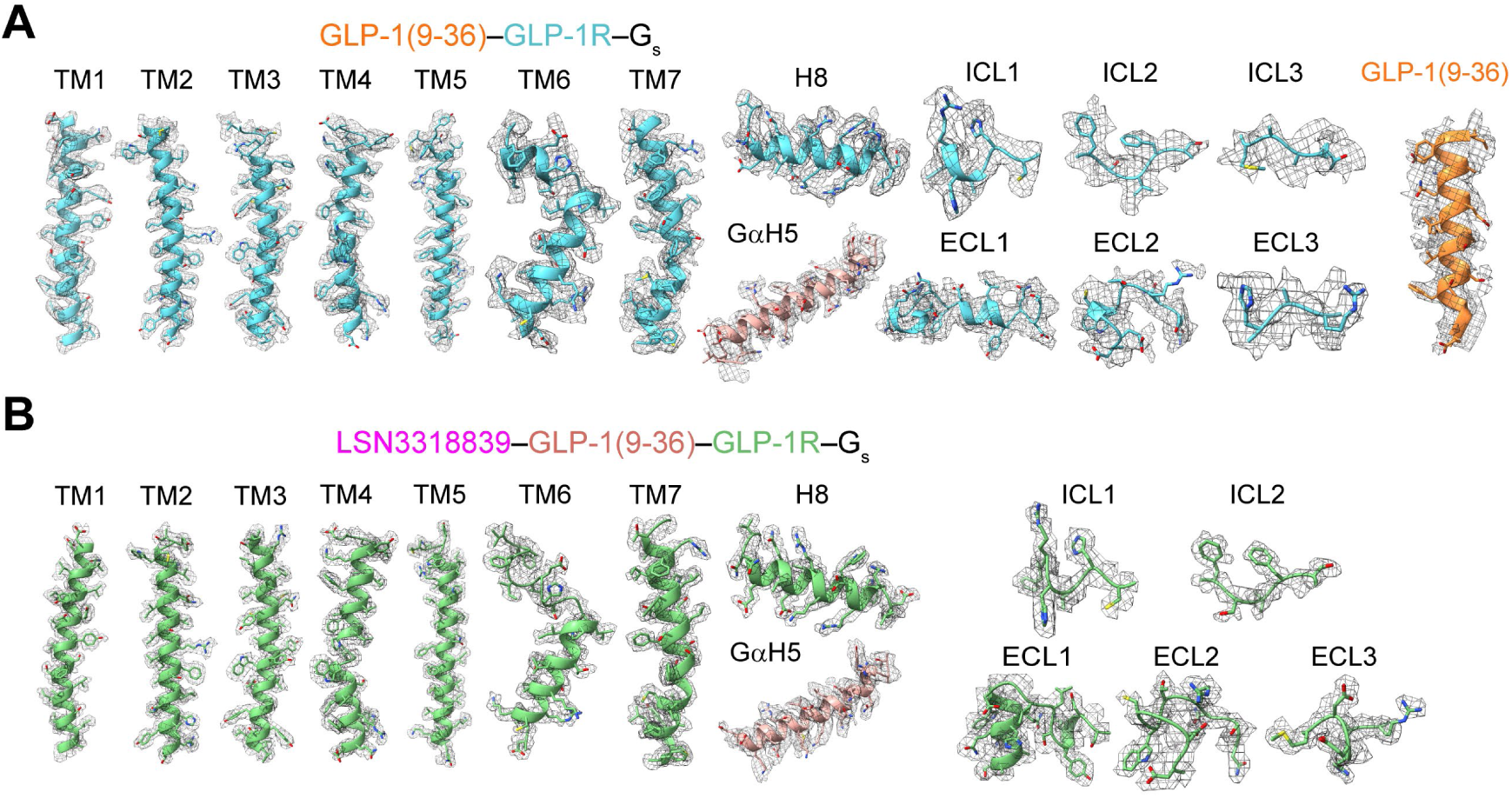
Near-atomic resolution models of the GLP-1(9-36)–GLP-1R–G_s_ and LSN3318839–GLP-1(9-36)–GLP-1R–G_s_ complexes in the cryo-EM density maps. (A) EM density map and model of the GLP-1(9-36)–GLP-1R–G_s_ complex are shown for all seven transmembrane α-helices (TMs 1-7), helix 8 (H8), intracellular loops 1-3 (ICL1–ICL3), extracellular loops 1-3 (ECL1–ECL3), the α5-helix of the Gα_s_ (GαH5) and GLP-1(9-36). (B) EM density map and model of the LSN3318839–GLP-1(9-36)–GLP-1R–G_s_ complexes are shown for TMs 1-7, H8, ICL1, ICL2, ECL1–ECL3 and GαH5.

**Figure S4.**
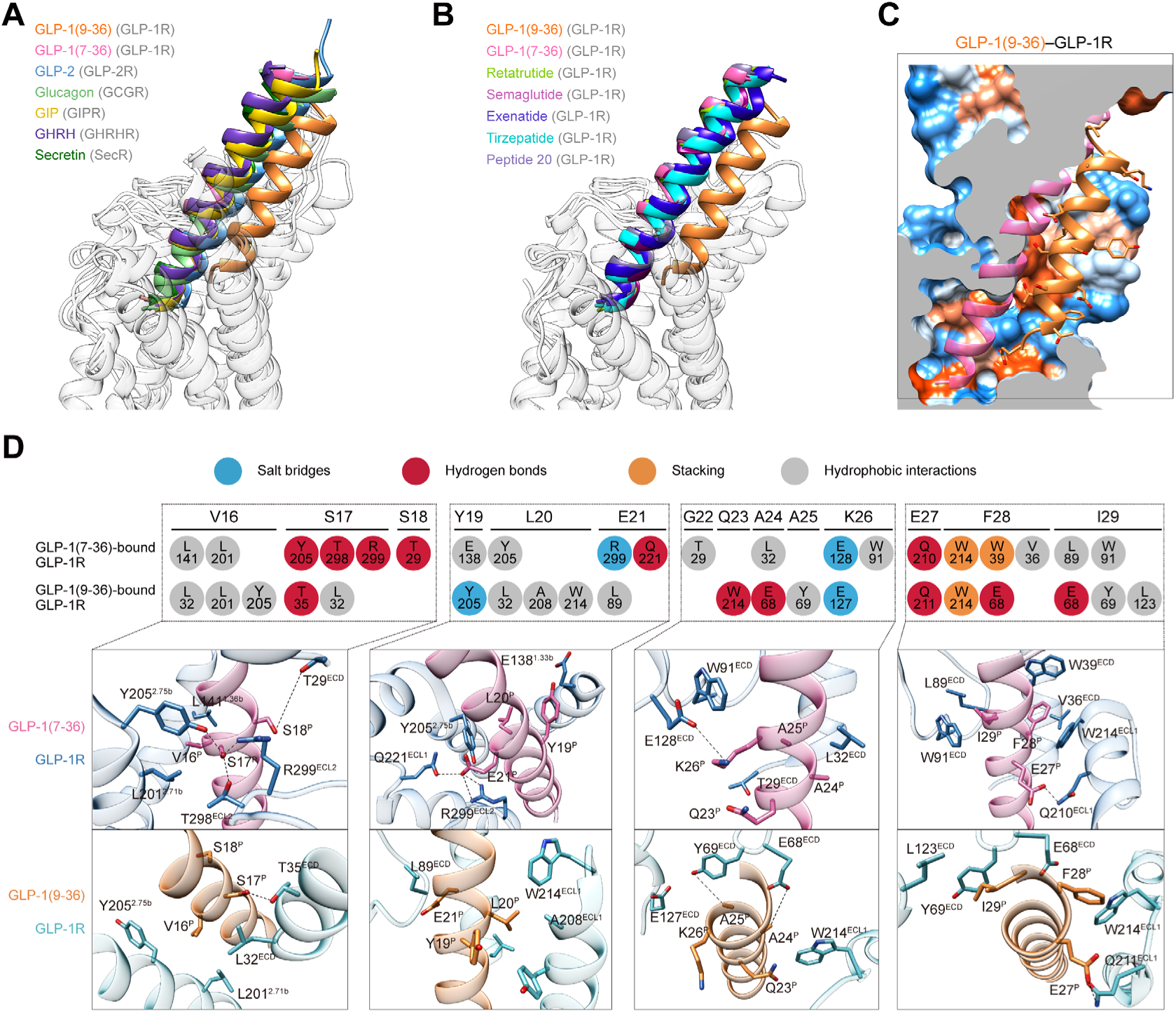
Molecular recognition of GLP-1(9-36) by GLP-1R. **(A) Comparison of hormone binding conformations in the glucagon receptor family.** (B) Comparison of peptide binding conformations in GLP-1R. The receptors are shown in transparent for clarity. (C) Comparison of GLP-1R interactions with the C-terminal halves of GLP-1(9-36) and GLP-1(7-36), described by fingerprint strings encoding different interaction types of the surrounding residues. Close-up views of these interactions are shown with polar contacts depicted as black dashed lines. Receptor residues are labeled with class B1 GPCR numbering and colored by interaction type: sky blue (salt bridge), red (hydrogen bond), orange (stacking) and gray (hydrophobic interactions).

**Figure S5.**
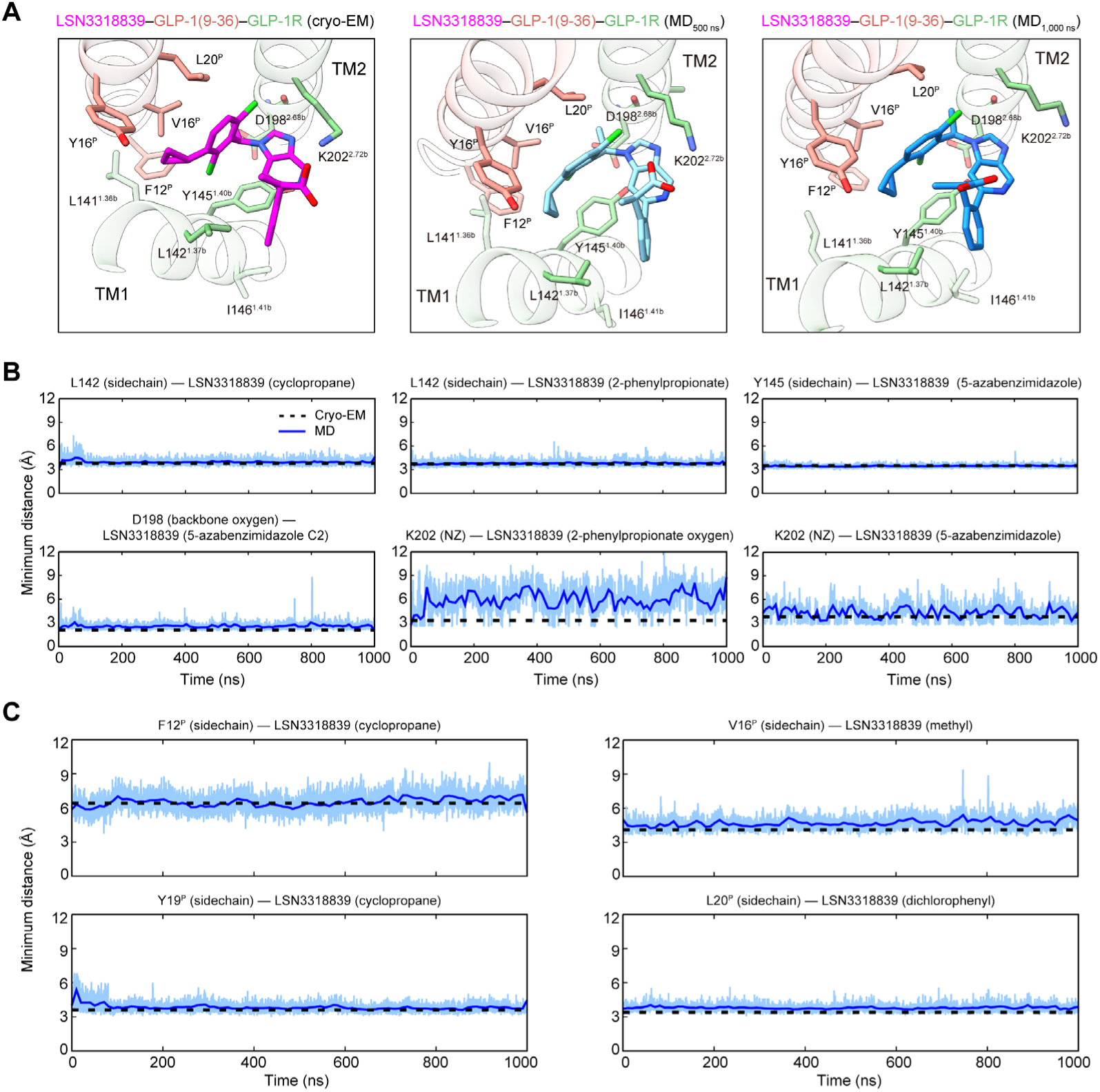
Molecular dynamics (MD) simulation of the binding pose of LSN3318839 at GLP-1(9-36)–GLP-1R. (A) Comparison of the LSN3318839 binding pose in the cryo-EM structure (left), MD snapshot at 500 ns (cyan) and the final snapshot at 1,000 ns (right). The pocket residues interacting with LSN3318839 are shown as sticks. (B) Representative minimum distances between the non-hydrogen atoms of LSN3318839 and the surrounding pocket residues from GLP-1R. (C) Representative minimum distances between the non-hydrogen atoms of LSN3318839 and the surrounding pocket residues from GLP-1(9-36). The thick and thin traces represent moving averages and original, unsmoothed values, respectively.

**Figure S6.**
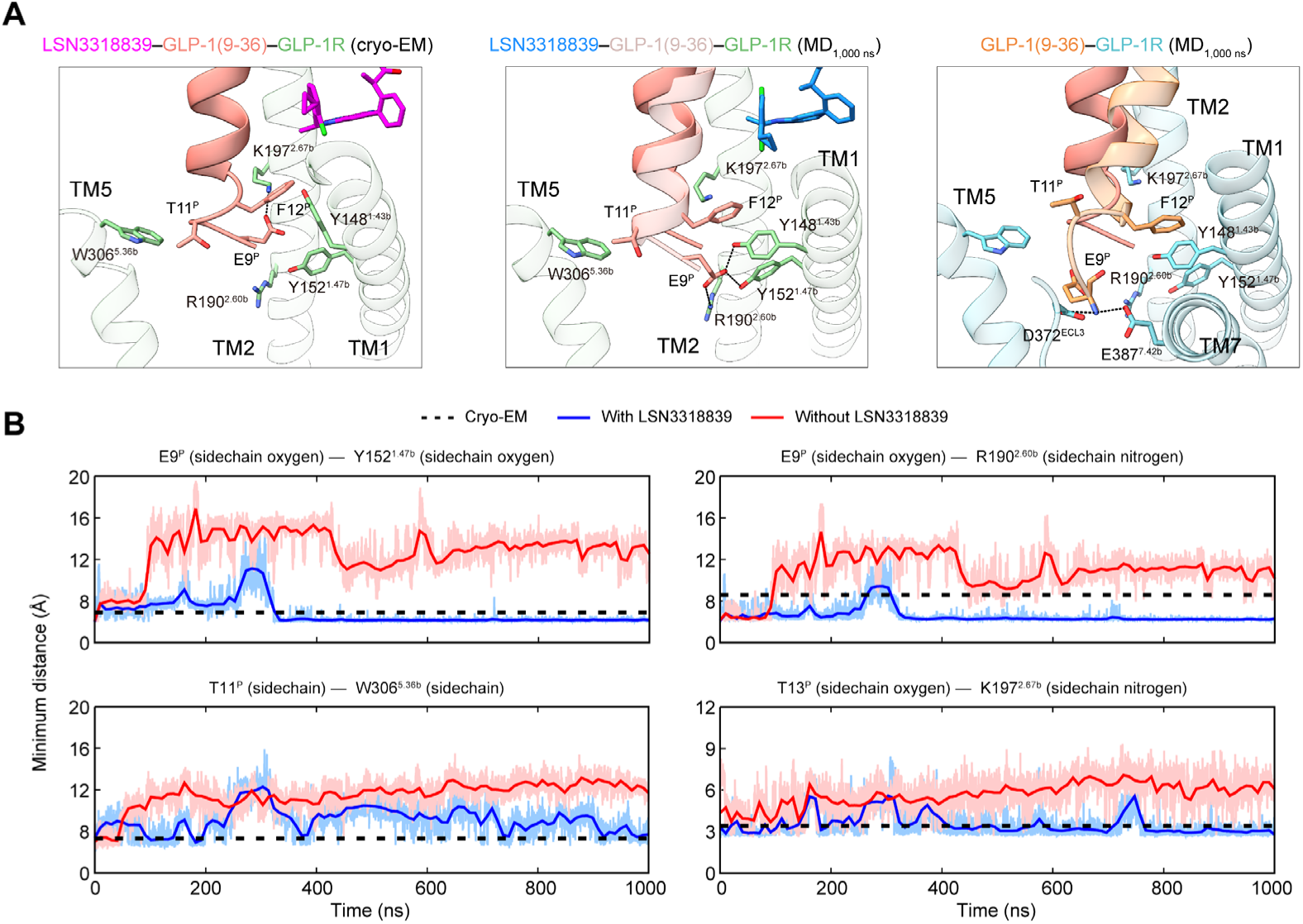
MD simulation of the GLP-1(9-36)–GLP-1R complex in the absence of LSN3318839. (A) Conformational comparison of the N terminus of GLP-1(9-36) in the cryo-EM structure (left), the final snapshots at 1,000 ns of the MD simulation of the LSN3318839– GLP-1(9-36)–GLP-1R complex (middle) and the GLP-1(9-36)–GLP-1R complex without LSN3318839 (right). The pocket residues interacting with the N terminus of GLP-1(9-36) are shown as sticks. (B) Representative minimum distances between the non-hydrogen atoms of the N terminus of GLP-1(9-36) and the surrounding pocket residues. The thick and thin traces represent moving averages and original, unsmoothed values, respectively.

**Table S1.** Cryo-EM data collection, model refinement and validation statistics.

|  | GLP-1(9-36)–<br>GLP-1R–G<sub>s</sub> | LSN3318839–GLP-1(9-36)–GLP-1R–G<sub>s</sub> |
| --- | --- | --- |
| **Data collection and processing** |  |  |
| Magnification | 81,000 | 81,000 |
| Voltage (kV) | 300 | 300 |
| Electron exposure (e-/Å<sup>2</sup>) | 80 | 80 |
| Defocus range (μm) | -1.2 to -2.2 | -1.2 to -2.2 |
| Pixel size (Å) | 1.071 | 1.071 |
| Symmetry imposed | C1 | C1 |
| Initial particle images (no.) | 12,688,111 | 6,146,299 |
| Final particle images (no.) | 99,120 | 238,182 |
| Map resolution (Å) | 3.0 | 3.2 |
| FSC threshold | 0.143 | 0.143 |
| Map resolution range (Å) | 3.0-3.6 | 3.2-4.1 |
| Map sharpening *B* factor (Å<sup>2</sup>) | –95 | –107 |
| **Refinement** |  |  |
| Initial model used (PDB code) | 6X18 | 6VCB |
| Model resolution (Å) | 3.2 | 4.1 |
| FSC threshold | 0.5 | 0.5 |
| Model resolution range (Å) | 2.9-5.0 | 3.2-5.0 |
| Model composition |  |  |
| Non-hydrogen atoms | 9,328 | 9,393 |
| Protein residues | 1,170 | 1,177 |
| *B* factors (Å<sup>2</sup>) |  |  |
| Protein | 106.68 | 84.31 |
| Ligand | – | 20 |
| Validation |  |  |
| MolProbity score | 2.37 | 2.18 |
| Clashscore | 7.77 | 9.24 |
| Poor rotamers (%) | 5.97 | 5.87 |
| Ramachandran plot |  |  |
| Favored (%) | 94.80 | 94.91 |
| Allowed (%) | 5.03 | 4.83 |
| Disallowed (%) | 0.17 | 0.26 |

**Table S2.** Interaction between peptide and GLP-1R.

| GLP-1 | GLP-1(7-36)–GLP-1R (6X18) | GLP-1(9-36)–GLP-1R | LSN3318839–GLP-1(9-36)–GLP-1R |
| --- | --- | --- | --- |
| H7<sup>P</sup> | Hydrogen bond with Q234<sup>3.37b</sup><br>Stacking with W306<sup>5.36b</sup><br>Hydrophobic contacts with V237<sup>3.40b</sup> | - | - |
| A8<sup>P</sup> | Hydrophobic contacts with L384<sup>7.39b</sup>, E387<sup>7.42b</sup> and L388<sup>7.43b</sup> | - | - |
| E9<sup>P</sup> | Salt bridge with R190<sup>2.60b</sup><br>Hydrogen bond with Y152<sup>1.47b</sup><br>Hydrophobic contacts with L388<sup>7.43b</sup> | Salt bridge with K197<sup>2.67b</sup><br>Hydrogen bond with Y148<sup>1.43b</sup><br>Stacking with Y148<sup>1.43b</sup> | Salt bridge with R190<sup>2.60b</sup> and K197<sup>2.67b</sup><br>Hydrogen bond with Y152<sup>1.47b</sup><br>Hydrophobic contacts with V237<sup>3.40b</sup> |
| G10<sup>P</sup> | Hydrophobic contacts with M233<sup>3.36b</sup> and W306<sup>5.36b</sup> | - | Hydrophobic contacts with M233<sup>3.36b</sup> and W306<sup>5.36b</sup> |
| T11<sup>P</sup> | Hydrogen bond with D372<sup>ECL3</sup><br>Hydrophobic contacts with L384<sup>7.39b</sup> | Hydrogen bond with L141<sup>1.36b</sup><br>Hydrophobic contacts with Y145<sup>1.40b</sup> | Hydrogen bond with D372<sup>ECL3</sup><br>Hydrophobic contacts with L384<sup>7.39b</sup> |
| F12<sup>P</sup> | Stacking with Y148<sup>1.43b</sup><br>Hydrophobic contacts with L141<sup>1.36b</sup>, L144<sup>1.39b</sup> and L388<sup>7.43b</sup> | Stacking with Y145<sup>1.40b</sup><br>Aromatic hydrogen bond with D198<sup>2.68b</sup><br>Hydrophobic contacts with K202<sup>2.72b</sup> | Stacking with Y148<sup>1.43b</sup><br>Hydrophobic contacts with L141<sup>1.36b</sup>, L144<sup>1.39b</sup> and L388<sup>7.43b</sup> |
| T13<sup>P</sup> | Hydrogen bond with K197<sup>2.67b</sup><br>Hydrophobic contacts with L201<sup>2.71b</sup> | Hydrogen bond with S31<sup>ECD</sup> and L32<sup>ECD</sup><br>Hydrophobic contacts with L201<sup>2.71b</sup> | Hydrogen bond with K197<sup>2.67b</sup><br>Hydrophobic contacts with L201<sup>2.71b</sup> |
| S14<sup>P</sup> | Hydrogen bond with N300<sup>ECL2</sup> (backbone) | - | Hydrogen bond with N300<sup>ECL2</sup> (backbone) |
| D15<sup>P</sup> | Salt bridge with R380<sup>7.35b</sup> | - | Salt bridge with R380<sup>7.35b</sup> |
| V16<sup>P</sup> | Hydrophobic contacts with L141<sup>1.36b</sup> and L201<sup>2.71b</sup> | Hydrophobic contacts with L32<sup>ECD</sup>, L201<sup>2.71b</sup> and Y205<sup>2.75b</sup> | Hydrophobic contacts with L141<sup>1.36b</sup> and L201<sup>2.71b</sup> |
| S17<sup>P</sup> | Hydrogen bond with Y205<sup>2.75b</sup>, T298<sup>45.52b</sup> and R299<sup>ECL2</sup> | Hydrogen bond with T35<sup>ECD</sup><br>Hydrophobic contacts with L32<sup>ECD</sup> | Hydrogen bond with Y205<sup>2.75b</sup>, T298<sup>45.52b</sup> and R299<sup>ECL2</sup> |
| S18<sup>P</sup> | Hydrogen bond with T29<sup>ECD</sup> | - | Hydrogen bond with T29<sup>ECD</sup> |
| Y19<sup>P</sup> | Hydrophobic contacts with E138<sup>1.33b</sup> | Stacking with Y205<sup>2.75b</sup> | Hydrogen bond with E138<sup>1.33b</sup> |
| L20<sup>P</sup> | Hydrophobic contacts with Y205<sup>2.75b</sup> | Hydrophobic contacts with L32<sup>ECD</sup>, | Hydrophobic contacts with Y205<sup>2.75b</sup> |
|  |  | A208<sup>ECL1</sup> and W214<sup>ECL1</sup> |  |
| E21<sup>P</sup> | Salt bridge with R299<sup>ECL2</sup><br>Hydrogen bond with Q221<sup>ECL1</sup> | Hydrophobic contacts with L89<sup>ECD</sup> | Salt bridge with R299<sup>ECL2</sup><br>Hydrogen bonds with S31<sup>ECD</sup>, L32<sup>ECD</sup><br>(backbone) and Y205 |
| G22<sup>P</sup> | Hydrophobic contacts with T29<sup>ECD</sup> | - | Hydrophobic contacts with T29<sup>ECD</sup> |
| Q23<sup>P</sup> | - | Aromatic hydrogen bond with W214<sup>ECL1</sup> | - |
| A24<sup>P</sup> | Hydrophobic contacts with L32<sup>ECD</sup> | Hydrogen bond with E68<sup>ECD</sup> | Hydrogen bond with Q210<sup>ECL1</sup><br>Hydrophobic contacts with L32<sup>ECD</sup> |
| A25<sup>P</sup> | - | Hydrophobic contacts with Y69<sup>ECD</sup> | - |
| K26<sup>P</sup> | Salt bridge with E128<sup>ECD</sup><br>Hydrophobic contacts with W91<sup>ECD</sup> |  | Salt bridge with E128<sup>ECD</sup><br>Hydrophobic contacts with W91<sup>ECD</sup> |
| E27<sup>P</sup> | Hydrogen bond with Q210<sup>ECL1</sup> | Hydrogen bond with Q211<sup>ECL1</sup> | Hydrophobic contacts with Q210<sup>ECL1</sup> |
| F28<sup>P</sup> | Stacking with W39<sup>ECD</sup> and W214<sup>ECL1</sup><br>Hydrophobic contacts with V36<sup>ECD</sup> | Stacking with W214<sup>ECL1</sup><br>Aromatic hydrogen bond with E68<sup>ECD</sup> | Stacking with W39<sup>ECD</sup> and W214<sup>ECL1</sup><br>Hydrophobic contacts with V36<sup>ECD</sup> |
| I29<sup>P</sup> | Hydrophobic contacts with L89<sup>ECD</sup> and W91<sup>ECD</sup> | Hydrogen bond with E68<sup>ECD</sup><br>Hydrophobic contacts with Y69<sup>ECD</sup> and<br>L123<sup>ECD</sup> | Hydrophobic contacts with L89<sup>ECD</sup> and<br>W91<sup>ECD</sup> |
| A30<sup>P</sup> | - | - | - |
| W31<sup>P</sup> | Stacking with W214<sup>ECL1</sup><br>Hydrogen bond with D215<sup>ECL1</sup><br>Hydrophobic contacts with W39<sup>ECD</sup> | - | Stacking with W214<sup>ECL1</sup><br>Hydrophobic contacts with W39<sup>ECD</sup> |
| L32<sup>P</sup> | Hydrophobic contacts with W39<sup>ECD</sup> and E68<sup>ECD</sup> | - | Hydrogen bond with R121<sup>ECD</sup><br>Hydrophobic contacts with W39<sup>ECD</sup> |
| V33<sup>P</sup> | Hydrogen bond with R121<sup>ECD</sup><br>Hydrophobic contacts with Y69<sup>ECD</sup> | - | Hydrogen bond with R121<sup>ECD</sup><br>Hydrophobic contacts with Y69<sup>ECD</sup> |
| K34<sup>P</sup> | - | - | - |
| G35<sup>P</sup> | - | - | - |
| R36<sup>P</sup> | Salt bridge with E68<sup>ECD</sup> | - | Salt bridge with E68<sup>ECD</sup> |

**Table S3.** Effect of GLP-1R mutation on peptide efficacy on cAMP signaling.

| Mutation | pEC<sub>50</sub> (M) of GLP-1(9-36) |  |  |  | pEC<sub>50</sub> (M) of GLP-1(7-36) |  |  |  |
| --- | --- | --- | --- | --- | --- | --- | --- | --- |
|  | + 0 nM<br>LSN3318839 | + 50 nM<br>LSN3318839 | + 500 nM<br>LSN3318839 | + 5,000 nM<br>LSN3318839 | + 0 nM<br>LSN3318839 | + 50 nM<br>LSN3318839 | + 500 nM<br>LSN3318839 | + 5,000 nM<br>LSN3318839 |
| WT | 6.44 ± 0.05 | 7.63 ± 0.08 | 8.48 ± 0.07 | 9.39 ± 0.08 | 10.79 ± 0.09 | 10.71 ± 0.08 | 10.64 ± 0.09 | 10.69 ± 0.11 |
| L142<sup>1.37b</sup>A | 6.22 ± 0.12 | 6.90 ± 0.06 | 7.62 ± 0.05 | 8.52 ± 0.08 | 10.84 ± 0.09 | 10.72 ± 0.10 | 10.71 ± 0.09 | 10.72 ± 0.10 |
| L142<sup>1.37b</sup>F | 6.56 ± 0.08 | 6.87 ± 0.05 | 7.48 ± 0.07 | 8.28 ± 0.08 | 10.85 ± 0.10 | 10.82 ± 0.09 | 10.68 ± 0.10 | 10.70 ± 0.12 |
| Y145<sup>1.40b</sup>A | 6.23 ± 0.07 | 6.93 ± 0.05 | 7.70 ± 0.04 | 8.50 ± 0.06 | 10.72 ± 0.08 | 10.61 ± 0.06 | 10.59 ± 0.08 | 10.61 ± 0.10 |
| D198<sup>2.68b</sup>A | N.A. | N.A. | N.A. | N.A. | 8.23 ± 0.09 | 8.54 ± 0.09 | 9.30 ± 0.09 | 10.16 ± 0.10 |
| K202<sup>2.72b</sup>A | N.A. | 6.51 ± 0.08 | 7.42 ± 0.06 | 8.33 ± 0.05 | 10.44 ± 0.07 | 10.45 ± 0.06 | 10.46 ± 0.08 | 10.48 ± 0.08 |
| Q234<sup>3.37b</sup>A | N.A. | 6.67 ± 0.09 | 7.45 ± 0.08 | 8.23 ± 0.10 | 9.60 ± 0.12 | 10.43 ± 0.10 | 10.52 ± 0.11 | 10.60 ± 0.12 |
| V237<sup>3.40b</sup>A | 6.45 ± 0.21 | 7.75 ± 0.08 | 8.60 ± 0.05 | 9.26 ± 0.08 | 10.43 ± 0.09 | 10.49 ± 0.10 | 10.47 ± 0.10 | 10.51 ± 0.12 |
| V237<sup>3.40b</sup>F | 6.40 ± 0.31 | 7.21 ± 0.05 | 8.06 ± 0.03 | 8.63 ± 0.06 | 8.38 ± 0.06 | 9.71 ± 0.05 | 10.45 ± 0.08 | 10.65 ± 0.05 |
| V237<sup>3.40b</sup>L | 6.57 ± 0.52 | 6.77 ± 0.06 | 7.62 ± 0.06 | 8.47 ± 0.06 | 10.01 ± 0.04 | 10.78 ± 0.04 | 10.95 ± 0.04 | 10.91 ± 0.05 |
| L388<sup>7.43b</sup>A | N.A. | 7.03 ± 0.05 | 7.96 ± 0.04 | 8.75 ± 0.04 | 9.36 ± 0.11 | 10.73 ± 0.05 | 10.96 ± 0.06 | 10.87 ± 0.07 |
| L388<sup>7.43b</sup>I | N.A. | 7.37 ± 0.07 | 8.27 ± 0.04 | 8.89 ± 0.05 | 10.62 ± 0.07 | 10.73 ± 0.05 | 10.66 ± 0.07 | 10.65 ± 0.06 |

GLP-1(9-36)/GLP-1(7-36)-induced cAMP production was measured in the presence of 0, 50, 500, or 5,000 nM LSN3318839. cAMP levels were determined by normalizing to the maximum response of GLP-1(7-36) at wildtype (WT) GLP-1R. Concentration-response curves were analyzed using a three-parameter logistic equation to obtain pEC<sub>50</sub> values. The experiments were carried out independently at least three times. Data shown are means ± S.E.M. N.A., not active.

**Table S4.** Effect of GLP-1R mutation on peptide potency displayed in cAMP accumulation.

| Mutation | E<sub>max</sub> (%) of GLP-1(9-36) |  |  |  | E<sub>max</sub> (%) of GLP-1(7-36) |  |  |  |
| --- | --- | --- | --- | --- | --- | --- | --- | --- |
|  | + 0 nM | + 50 nM | + 500 nM | + 5,000 nM | + 0 nM | + 50 nM | + 500 nM | + 5,000 nM |
|  | LSN3318839 | LSN3318839 | LSN3318839 | LSN3318839 | LSN3318839 | LSN3318839 | LSN3318839 | LSN3318839 |
| WT | 84.77 ± 2.11 | 97.94 ± 2.50 | 97.96 ± 1.99 | 96.85 ± 2.01 | 100.00 ± 2.66 | 100.21 ± 2.38 | 100.01 ± 2.51 | 99.76 ± 3.10 |
| L142<sup>1.37b</sup>A | 51.17 ± 3.06 | 94.12 ± 2.37 | 98.91 ± 1.55 | 98.42 ± 2.25 | 101.11 ± 2.65 | 100.50 ± 2.92 | 100.14 ± 2.68 | 100.01 ± 2.78 |
| L142<sup>1.37b</sup>F | 83.77 ± 2.97 | 91.78 ± 2.04 | 97.53 ± 2.43 | 97.38 ± 2.31 | 100.73 ± 2.76 | 99.93 ± 2.52 | 100.10 ± 2.75 | 100.02 ± 3.30 |
| Y145<sup>1.40b</sup>A | 61.54 ± 2.28 | 90.38 ± 1.72 | 95.00 ± 1.34 | 93.29 ± 1.49 | 100.14 ± 2.13 | 100.05 ± 1.78 | 98.99 ± 2.15 | 99.37 ± 2.74 |
| D198<sup>2.68b</sup>A | N.A. | N.A. | N.A. | N.A. | 100.44 ± 4.63 | 102.91 ± 3.85 | 99.85 ± 3.39 | 99.99 ± 3.24 |
| K202<sup>2.72b</sup>A | N.A. | 78.55 ± 2.58 | 97.57 ± 1.91 | 98.40 ± 1.34 | 101.95 ± 2.02 | 101.39 ± 1.81 | 101.62 ± 2.30 | 101.49 ± 2.55 |
| Q234<sup>3.37b</sup>A | N.A. | 72.82 ± 2.90 | 89.24 ± 2.61 | 88.88 ± 2.97 | 98.08 ± 3.99 | 98.17 ± 3.02 | 97.79 ± 3.01 | 97.91 ± 3.31 |
| V237<sup>3.40b</sup>A | 52.69 ± 5.32 | 90.68 ± 2.58 | 89.82 ± 1.36 | 89.74 ± 1.87 | 96.71 ± 2.61 | 96.06 ± 2.70 | 94.53 ± 2.70 | 94.55 ± 3.37 |
| V237<sup>3.40b</sup>F | 17.76 ± 2.43 | 92.37 ± 1.90 | 96.39 ± 1.10 | 97.53 ± 1.80 | 102.14 ± 2.60 | 100.00 ± 1.74 | 100.48 ± 2.24 | 100.33 ± 1.34 |
| V237<sup>3.40b</sup>L | 10.25 ± 1.36 | 92.83 ± 2.37 | 99.4 ± 1.95 | 98.86 ± 1.66 | 101.00 ± 1.31 | 100.69 ± 1.12 | 100.47 ± 1.18 | 100.81 ± 1.48 |
| L388<sup>7.43b</sup>A | N.A. | 85.60 ± 1.66 | 96.40 ± 1.23 | 98.69 ± 1.10 | 104.12 ± 4.24 | 101.39 ± 1.51 | 100.58 ± 1.66 | 101.03 ± 1.91 |
| L388<sup>7.43b</sup>I | N.A. | 84.74 ± 2.15 | 93.91 ± 1.24 | 95.82 ± 1.21 | 100.84 ± 2.02 | 99.92 ± 1.39 | 99.61 ± 1.98 | 100.18 ± 1.63 |

GLP-1(9-36)/GLP-1(7-36)-induced cAMP production was measured in the presence of 0, 50, 500, or 5,000 nM LSN3318839. cAMP levels were determined by normalizing to the maximum response of GLP-1(7-36) at WT GLP-1R. Concentration-response curves were analyzed using a three-parameter logistic equation to obtain E<sub>max</sub> values. The experiments were carried out independently at least three times. Data shown are means ± S.E.M. N.A., not active.

